# A Reinforcement Learning Approach for Modeling Organic Compound-Induced Antimicrobial Resistance Dynamics

**DOI:** 10.64898/2025.12.18.695270

**Authors:** Hayden D. Hedman

**Affiliations:** School of Computer Science, Georgia Institute of Technology, Atlanta, GA, USA

**Keywords:** reinforcement learning, antimicrobial resistance, copper exposure, minimum inhibitory concentration (MIC), Q-learning, agent-based modeling, simulation framework

## Abstract

This study presented an exploratory reinforcement learning (RL)-based simulation framework for examining antimicrobial resistance (AMR) dynamics under repeated exposure to an organic antimicrobial stressor, using copper as a representative model compound. Within a simplified and explicitly constrained simulation environment, three agent strategies were evaluated: random action selection, a rule-based heuristic, and a tabular Q-learning agent. Simulations were conducted over fixed-length 40-cycle episodes in which agents adjusted copper exposure in response to evolving resistance-related state variables. Across experimental runs, the Q-learning agent exhibited lower cumulative antibiotic resistance burden, measured by the area under the curve (AUC) of minimum inhibitory concentration (MIC) values for chloramphenicol and polymyxin B, while also maintaining lower cumulative copper exposure relative to the rule-based and random baselines. The rule-based agent demonstrated intermediate performance, whereas the random agent showed higher variability and less stable resistance trajectories. These differences reflected divergence in simulated resistance dynamics over time rather than short-term fluctuations in resistance burden. Rather than providing predictive or mechanistic insight into microbial evolution, this work introduced an interpretable RL-based simulation framework intended to support comparative evaluation of sequential decision-making strategies under constrained observability, where high-resolution diagnostics or detailed biological measurements may be unavailable. Together, the results supported the use of reinforcement learning as a flexible methodological framework for studying AMR dynamics as a feedback-driven control problem under simplified and transparent assumptions.

## 1. Introduction

Antimicrobial resistance (AMR) continues to be one of the most critical global health challenges, with resistance to conventional antibiotics increasing at an alarming rate worldwide (*1–3*). The widespread emergence of resistant pathogens has undermined the long-term effectiveness of many frontline antimicrobial agents, placing increasing strain on clinical, agricultural, and public health systems (*4–6*). While antibiotic development remains an important priority, the pace of novel drug discovery has not kept up with the rate at which resistance evolves (*7,8*), motivating interest in complementary strategies for managing antimicrobial pressure rather than relying exclusively on new synthetic compounds (*9–11*).

Within this context, alternative and organic antimicrobial agents have attracted growing attention as potential tools for modulating resistance dynamics (*12–14*). Among these, copper has long been recognized for its broad-spectrum antimicrobial activity and its capacity to inhibit microbial growth through multiple, non-specific mechanisms (*15–18*). Copper and copper-based materials have been explored across diverse settings, including surface coatings, agricultural applications, and environmental controls, where they may reduce microbial burden without acting as direct therapeutic replacements for antibiotics (*15,16,18*). However, while copper shows consistent antimicrobial activity, evidence on its longer-term influence on AMR trajectories is mixed and remains incompletely characterized across settings and exposure regimes (*12,19,20*). In particular, the interaction between sublethal copper exposure and the evolution of antibiotic resistance poses unresolved questions, especially given evidence that metal stressors can contribute to co-selection and persistence of resistance phenotypes under certain conditions (*17,21*). Rather than serving as a substitute for conventional antimicrobial therapy, copper provides a biologically grounded model compound for examining how organic antimicrobial pressure influences resistance dynamics over time. Its use as a model stressor allows investigation of resistance evolution under repeated, variable exposure without the confounding assumptions associated with optimized therapeutic dosing. This perspective motivates the need for systematic and adaptive experimentation frameworks capable of capturing how resistance accumulates, stabilizes, or partially reverses across sequential exposure cycles.

The evolution of antimicrobial resistance can be naturally framed as a sequential decision-making and feedback-control problem (*22,23*). Resistance does not emerge from a single exposure event but instead reflects cumulative responses to repeated environmental pressures, where past decisions shape future susceptibility and growth outcomes. From this perspective, antimicrobial management involves a series of interdependent control decisions made under uncertainty, incomplete observability, and delayed feedback. Reinforcement learning (RL) provides a computational framework well suited to such settings, as it explicitly models sequential decision-making in dynamic environments where system dynamics may be stochastic or only partially known.

Recent foundational work has demonstrated that reinforcement learning can learn adaptive intervention strategies capable of influencing resistance trajectories in complex evolutionary systems defined by antibiotic fitness landscapes, even under noisy or delayed feedback conditions (*22,24*). These studies establish the feasibility of RL-based approaches for AMR control and illustrate how adaptive policies can outperform static treatment paradigms when resistance dynamics unfold over time. However, existing RL applications in AMR research have largely focused on synthetic antibiotics and highly structured treatment environments, often assuming detailed fitness landscapes, rich diagnostic information, or tightly controlled experimental conditions.

As a result, comparatively little work has examined how adaptive decision-making frameworks behave when applied to organic antimicrobial compounds or simplified settings that more closely reflect constrained clinical, agricultural, or infrastructural capacities. In many real-world environments, resistance management decisions must be made with limited diagnostic feedback, coarse measurements of susceptibility, and restricted ability to precisely modulate antimicrobial exposure. Understanding how learning-based and heuristic strategies perform under such constraints represents an important methodological gap.

Building on this foundation, the present study applies a reinforcement learning– based simulation framework using copper as a representative organic antimicrobial stressor to explore how different decision-making strategies respond to antimicrobial pressure across variable and constrained environments. The goal of this work is not to evaluate the clinical effectiveness of copper itself, but to examine how varying levels of adaptability influence resistance dynamics under simplified, interpretable assumptions. By abstracting resistance evolution using commonly reported microbiological indicators such as minimum inhibitory concentrations (MICs) and derived resistance burden indices, the framework emphasizes conceptual clarity and methodological transparency rather than biological or computational complexity.

To this end, three agent strategies are considered: (1) a random agent representing uncontrolled or unstructured exposure, (2) a rule-based agent implementing fixed decision criteria based on observed resistance trends, and (3) a Q-learning agent employing adaptive decision-making through iterative interaction with the environment. By comparing these agents within a shared simulation environment, this study provides an exploratory framework for understanding how adaptive control strategies may perform relative to static or heuristic approaches when resistance evolves over repeated exposure cycles.

Collectively, this work situates antimicrobial resistance within a systems and control-oriented perspective, highlighting the potential role of reinforcement learning as a methodological tool for studying AMR dynamics under uncertainty. Rather than offering prescriptive treatment recommendations, the framework introduced here aims to support future investigations into AI-assisted modeling approaches that bridge microbiology, systems thinking, and adaptive decision-making in the study of antimicrobial resistance.

## 2. Materials and Methods

### 2.1. Overview of Study and Framework

This study implemented a reinforcement learning (RL)–based simulation framework to model antibiotic resistance dynamics under varying copper exposure conditions across discrete experimental cycles. The experimental setup was rooted in copper concentration estimates reported in the foundational study by Li et al. (2021) (*21*), with parameters adapted to simulate copper exposure and antibiotic resistance evolution in *Escherichia coli K12 and Staphylococcus aureus*. The RL environment and agents were implemented in Python (Gymnasium, NumPy, and Matplotlib) for simulation, computation, and figure generation. Simulations modeled bacterial populations exposed to varying copper concentrations in combination with antibiotics chloramphenicol and polymyxin B (*21*). Three agents were evaluated: (1) a random policy agent, (2) a rule-based agent, and (3) a Q-learning agent. These models were designed to simulate control strategies for antimicrobial resistance dynamics under copper exposure.

Simulations were conducted over fixed-length episodes, each consisting of 40 cycles. At the start of each episode, the environment was reset to baseline conditions and the agent state was reinitialized. Within each cycle, agents selected a copper adjustment action based on the current state, and resistance- and growth-related outcomes were updated according to stochastic transition rules. Agent performance was evaluated by aggregating per-cycle outputs across repeated simulation runs. Simulations rely on pseudo-random number generation within the environment transition dynamics; repeated episodes were used to characterize variability in agent trajectories.

### 2.2. Reinforcement Learning Agent Design

The reinforcement learning (RL) agents interacted with a simulation environment designed to capture key features of copper-driven antimicrobial pressure and resistance dynamics over discrete experimental cycles (*21*). The environment represents bacterial populations exposed to variable copper concentrations in the presence of antibiotics, without explicitly modeling underlying molecular or genetic mechanisms.

The environment state was represented at each cycle as a five-dimensional vector consisting of current copper concentration (mg/L), minimum inhibitory concentration (MIC) values for chloramphenicol and polymyxin B, cycle count, and a normalized resistance burden index. These state variables were selected to reflect commonly reported microbiological indicators of resistance and antimicrobial pressure, including MIC values and derived resistance burden summaries (*25,26*). Copper concentration was bounded within an experimentally plausible range (0–100 mg/L), while MIC values were initialized at low baseline levels and allowed to evolve dynamically over time. The cycle count tracked progression within each episode, and the resistance burden index summarized the cumulative antibiotic resistance burden across antibiotics.

The action space consisted of three discrete control actions: decrease copper concentration, maintain the current level, or increase copper concentration. These actions corresponded to fixed stepwise adjustments in copper exposure applied at each cycle. Episodes were initialized at a sublethal copper concentration of 10 mg/L, consistent with exposure estimates reported by Li and colleagues (*21*), with copper levels constrained to remain within 0–100 mg/L throughout simulation.

Other biological dynamics, including changes in MIC values and growth inhibition, were modeled using stochastic update rules intended to represent resistance progression and partial reversibility under reduced antimicrobial pressure rather than explicit mechanistic mutation processes. To promote interpretability and reduce state-space complexity, continuous state variables were discretized into coarse bins prior to Q-learning. This design choice emphasized conceptual biological relevance and clarity over fine-grained mechanistic resolution, consistent with the exploratory objectives of the study.

### 2.3. Resistance Evolution

Consistent with prior work framing antimicrobial resistance as a feedback-control problem under noisy and partially observed dynamics (*22*), changes in antibiotic resistance were modeled using stochastic update rules intended to capture conceptual resistance progression under antimicrobial pressure rather than explicit molecular or genetic mechanisms. At each cycle, the probability of resistance amplification increased as a function of copper concentration, representing selection pressure imposed by the antimicrobial environment. When amplification occurred, minimum inhibitory concentration (MIC) values for chloramphenicol and polymyxin B were increased by antibiotic-specific stochastic increments.

To reflect partial phenotypic reversibility under reduced antimicrobial pressure, MIC values were allowed to decrease modestly when copper concentrations fell below a low-exposure threshold. These decreases were implemented stochastically and were smaller in magnitude than resistance increases, reflecting the asymmetric nature of resistance acquisition and loss. MIC values were constrained to non-negative values throughout simulation.

### 2.4. Resistance Burden Index

A resistance burden index was derived deterministically as a normalized summary of the combined antibiotic resistance burden, calculated as one minus the normalized sum of MIC values for chloramphenicol and polymyxin B. This index does not represent empirical growth inhibition, cell viability, or growth kinetics, but instead serves as an abstract, bounded proxy for aggregate resistance burden within the modeled population. Higher index values correspond to lower resistance burden, while lower values reflect elevated resistance. The index was included to support interpretability and reward shaping rather than to model biological growth processes. Because the index is a deterministic function of MIC values, it does not introduce independent state information beyond the resistance measures themselves. MIC values and derived indices were treated as abstract state variables representing resistance burden rather than direct predictors of microbial growth or clinical susceptibility

### 2.5. Reward Function

Agent learning was guided by a scalar reward function designed to balance resistance suppression and copper minimization. Rewards increased with lower aggregate resistance burden, as reflected by the resistance burden index, and were penalized by elevated MIC values and increased copper exposure. Weighting terms were selected to keep individual reward components on comparable scales and to discourage trivial strategies that rely on persistently high copper concentrations. Reward parameters were chosen to reflect qualitative trade-offs rather than to maximize performance through extensive tuning, consistent with the exploratory aims of the study.

### 2.6. Experiment Protocol and Data Generation

Experiments were conducted in a fixed-length episodic simulation setting. Each episode began with a reset of the environment to baseline conditions and proceeded for a maximum of 40 cycles. At each cycle, an agent selected one of three discrete copper-control actions (decrease, maintain, increase), after which the environment updated copper concentration, minimum inhibitory concentration (MIC) values for chloramphenicol and polymyxin B, the growth inhibition proxy, and reward. Per-cycle outcomes were recorded for each episode, including copper concentration, MIC trajectories for both antibiotics, resistance burden index values, and reward.

The random-policy and rule-based agents were evaluated over 50 independent episodes each. The Q-learning agent was run for 100 episodes with online temporal-difference updates applied at each cycle. The Q-learning agent reflects adaptive control through online learning, whereas the random and rule-based agents are non-learning baselines with fully specified policies.

### 2.7. Random Policy Agent

The random policy agent served as a negative-control baseline and selected copper-control actions uniformly at random at each cycle. This agent did not incorporate any information from the environment state and did not adapt its behavior based on observed resistance dynamics or prior outcomes. As such, the random policy represents an uncontrolled exposure scenario in which copper adjustments are applied without feedback or strategic intent.

Performance of the random policy reflects baseline resistance trajectories under non-optimized copper exposure and provides a reference point for evaluating the added value of structured and adaptive decision-making strategies.

### 2.8. Rule-Based Agent

The rule-based agent followed predefined decision rules to adjust copper exposure based on observed antibiotic resistance, as measured by MIC values. Copper levels were reduced when resistance increased, maintained when MIC values remained stable, and increased when resistance decreased. This agent reflects a protocol-driven decision strategy that could be applied in settings with limited computational or diagnostic resources. Its performance was compared with that of the Q-learning agent to evaluate the potential benefits of adaptive strategies for managing resistance dynamics.

### 2.9. Q-Learning Agent

The Q-learning agent implemented a tabular, model-free reinforcement learning approach to learn an adaptive copper-control policy through repeated interaction with the simulation environment. Because the environment state variables are continuous, the agent used coarse discretization to map each state to a finite set of bins prior to learning. Specifically, copper concentration (0–100 mg/L), MIC values for chloramphenicol and polymyxin B (0–512), and cycle count (0–max cycles) were each discretized into 10 equal-width bins and encoded as a four-dimensional discrete state index (*27*). The resistance burden index was retained for reward computation but was not included in the Q-table state representation.

Action selection followed an ε-greedy policy (ε = 0.10): with probability ε the agent sampled a random action, otherwise it selected the action with the highest estimated state–action value. Q-values were updated online at each cycle using the standard temporal-difference update rule with learning rate α = 0.10 and discount factor γ = 0.95. The Q-table persisted across episodes, allowing cumulative learning over repeated simulation runs.

### 2.10. Longitudinal Modeling of Resistance Trajectories

To quantify differences in antimicrobial resistance trajectories across agent strategies over repeated simulation cycles, longitudinal statistical models were applied to the per-cycle simulation outputs. In addition to summary performance metrics, including the area under the curve (AUC) of minimum inhibitory concentration (MIC) values, cumulative copper burden, and mean resistance burden index value, cumulative copper burden, and mean growth inhibition, these models were used to directly evaluate temporal patterns in resistance evolution across agents.

Resistance outcomes for chloramphenicol and polymyxin B were analyzed using generalized linear mixed models (GLMMs) implemented in R (packages: lme4, lmerTest, readr, and emmeans) (*28*). Mixed-effects modeling was selected to account for the repeated-measures structure (*29,30*) of the simulation data, in which MIC values were recorded across multiple cycles within each episode. This approach allows separation of systematic effects attributable to agent behavior from stochastic variability across simulation runs.

For each antibiotic, MIC values were modeled as a function of simulation cycle, agent type (random, rule-based, or Q-learning), and their interaction, with episode included as a random intercept to capture between-episode heterogeneity. The inclusion of cycle-by-agent interaction terms enabled direct comparison of resistance trajectories over time, rather than reliance on single time-point or aggregate measures alone.

Model results are reported as estimated effects with 95% confidence intervals (CIs), providing interpretable measures of uncertainty and effect magnitude. Pairwise contrasts between agent types were evaluated using marginal means with multiplicity-adjusted confidence intervals to assess meaningful differences in resistance dynamics across control strategies. Together, these analyses complement AUC-based summaries by providing a model-based assessment of how resistance trajectories diverge under different decision-making frameworks.

### 2.11. Reproducibility and Generative AI Statement

All code used for experiments, data analysis, and figure generation is provided in the supplementary material (SM1). Scripts are organized with clear instructions for setup and execution. Raw data and generated figures are saved in CSV and image formats, respectively, allowing validation and extension of the framework. Generative AI tools were used for coding assistance, troubleshooting, and initial idea generation; all outputs were reviewed prior to inclusion in the final manuscript.

## 3. Results

### 3.1. Agent-level resistance dynamics based on AUC metrics

Across both antibiotics, the Q-learning agent exhibited lower cumulative resistance exposure than the random and rule-based agents, as measured by the area under the curve (AUC) of minimum inhibitory concentration (MIC) values (Table 1; Table 2; Figures 1–4). Bootstrap contrasts indicated lower AUC MIC values for the Q-learning agent relative to the random agent for both chloramphenicol and polymyxin B. Differences between the Q-learning and rule-based agents were smaller, consistent with intermediate performance under rule-based decision-making (Table 1).

**Table 1.**
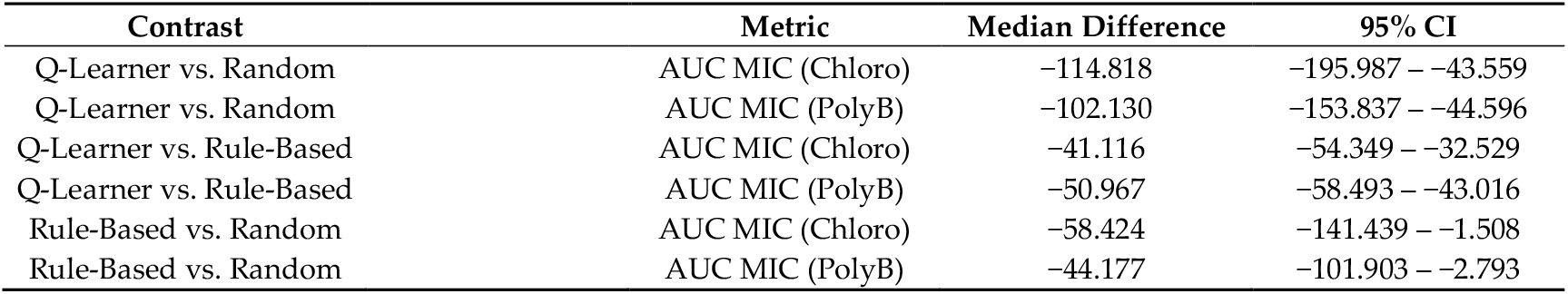
Bootstrap AUC summary for agent comparisons.

**Table 2.** Summary metrics for agent performance (Mean and Standard Deviation) for Chloramphenicol (Chloro) and Polymyxin B (PolyB).

| Agent | AUC MIC (Chloro)<br>Mean (SD) | AUC MIC (PolyB)<br>Mean (SD) | Copper Burden<br>Mean (SD) | Mean Growth<br>Inhibition<br>Mean (SD) |
| --- | --- | --- | --- | --- |
| Q-Learner | 10.131 (8.917) | 22.229 (12.035) | 15.050 (9.087) | 0.999 (0.000) |
| Random | 183.340 (175.391) | 149.517 (113.335) | 564.200 (394.399) | 0.992 (0.007) |
| Rule-Based | 69.739 (52.945) | 76.042 (32.698) | 142.200 (91.390) | 0.996 (0.002) |

**Table 3.** Changes in antibiotic minimum inhibitory concentration (MIC) across simulation cycles by agent type, with 95% confidence intervals.

| Effect | Chloramphenicol Estimate (95% CI) | Polymyxin B Estimate (95% CI) |
| --- | --- | --- |
| Cycle | −0.031 (−0.038 – −0.024) | −0.051 (−0.055 – −0.046) |
| Random | 0.062 (−0.229 – 0.352) | −0.222 (−0.412 – −0.033) |
| Rule-Based | 1.879 (1.589 – 2.169) | 1.113 (0.924 – 1.302) |
| Cycle × Random | 0.214 (0.202 – 0.226) | 0.170 (0.162 – 0.178) |
| Cycle × Rule-Based | −0.018 (−0.030 – −0.006) | 0.012 (0.004 – 0.020) |

**Figure 1.**
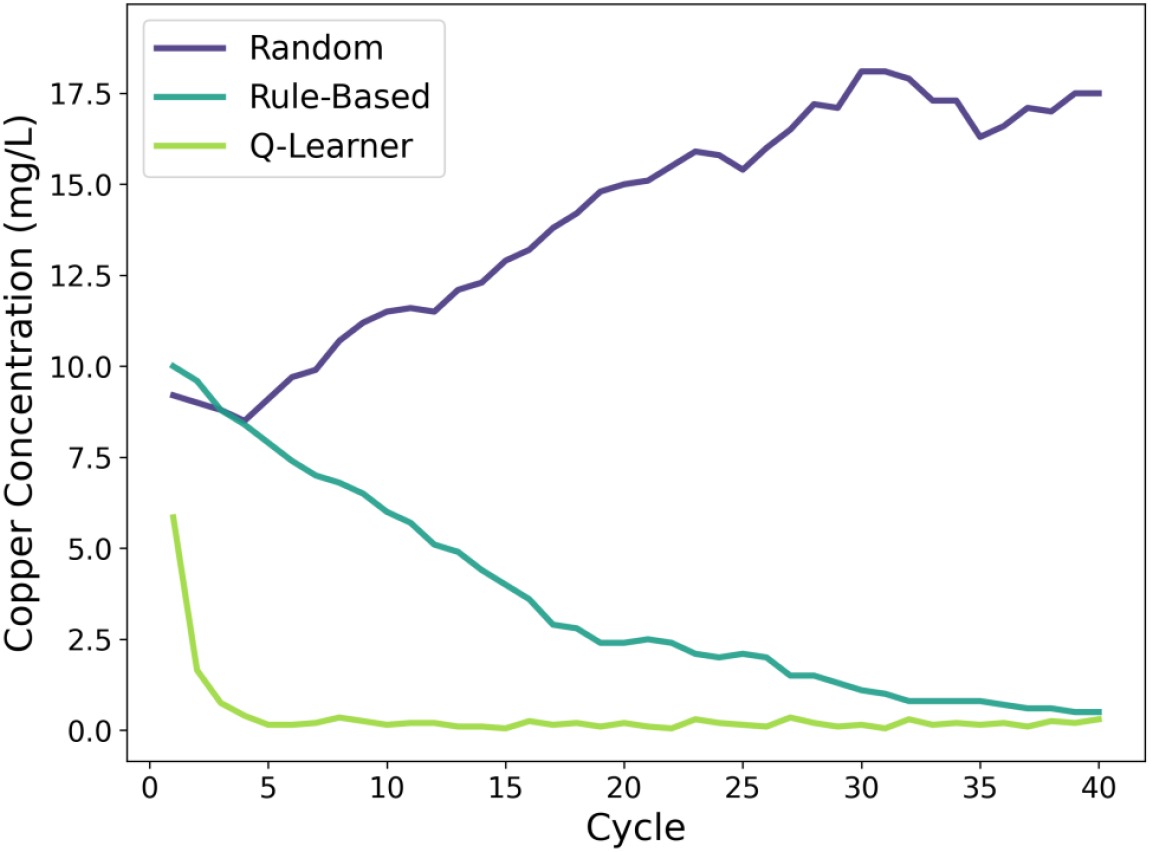
Copper concentration (mg/L) across experimental cycles for each agent. The figure shows how copper exposure was adjusted over cycles by the random, rule-based, and Q-learning agents.

**Figure 2.**
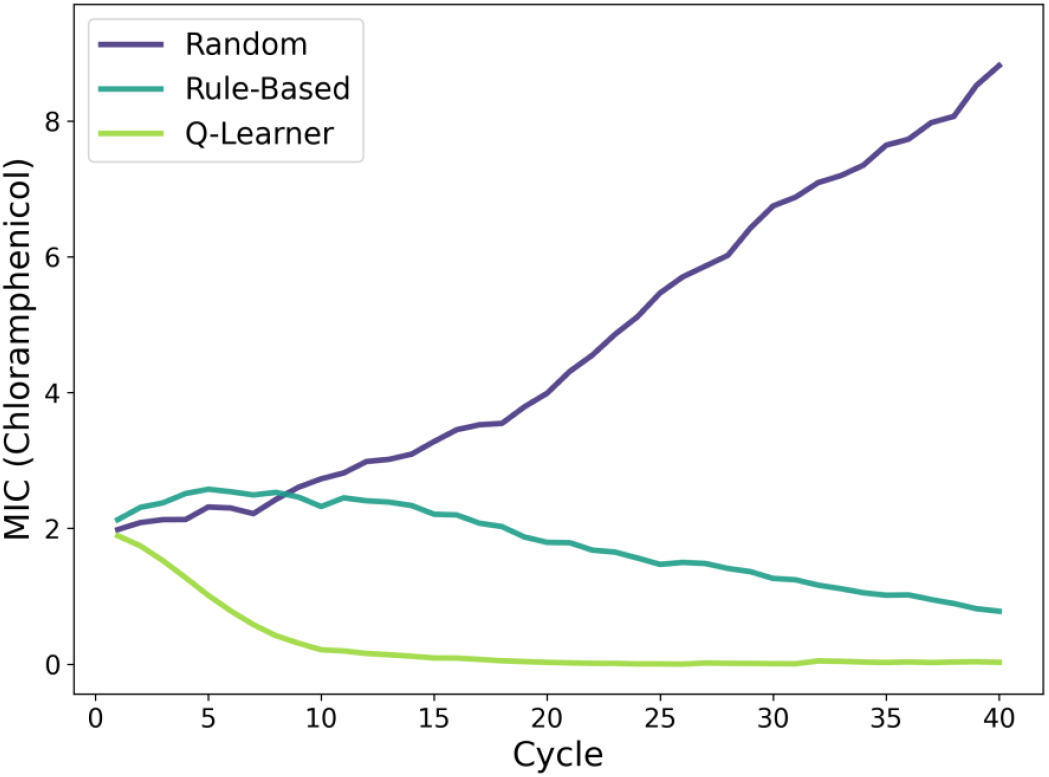
Chloramphenicol MIC trajectories across experimental cycles, illustrating differences in agent-driven control of resistance.

**Figure 3.**
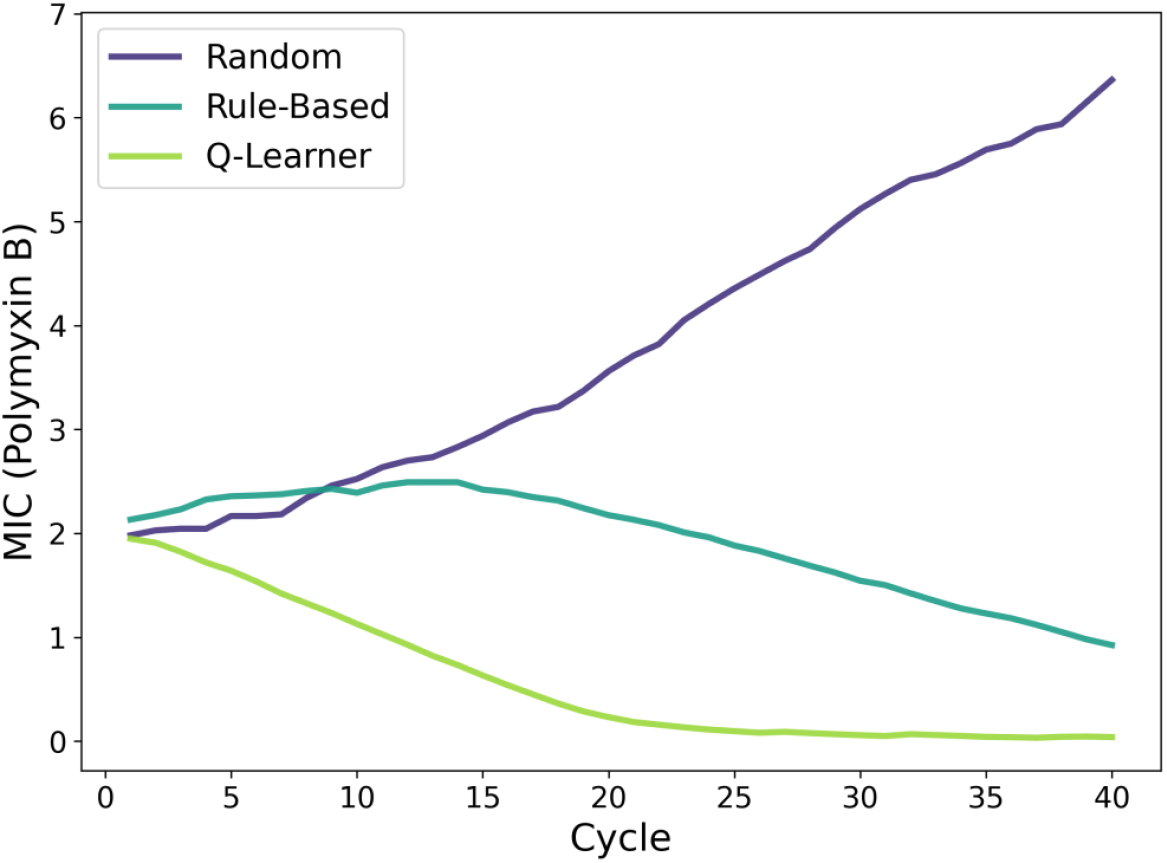
Polymyxin B minimum inhibitory concentration (MIC) across experimental cycles for the random, rule-based, and Q-learning agents.

**Figure 4.**
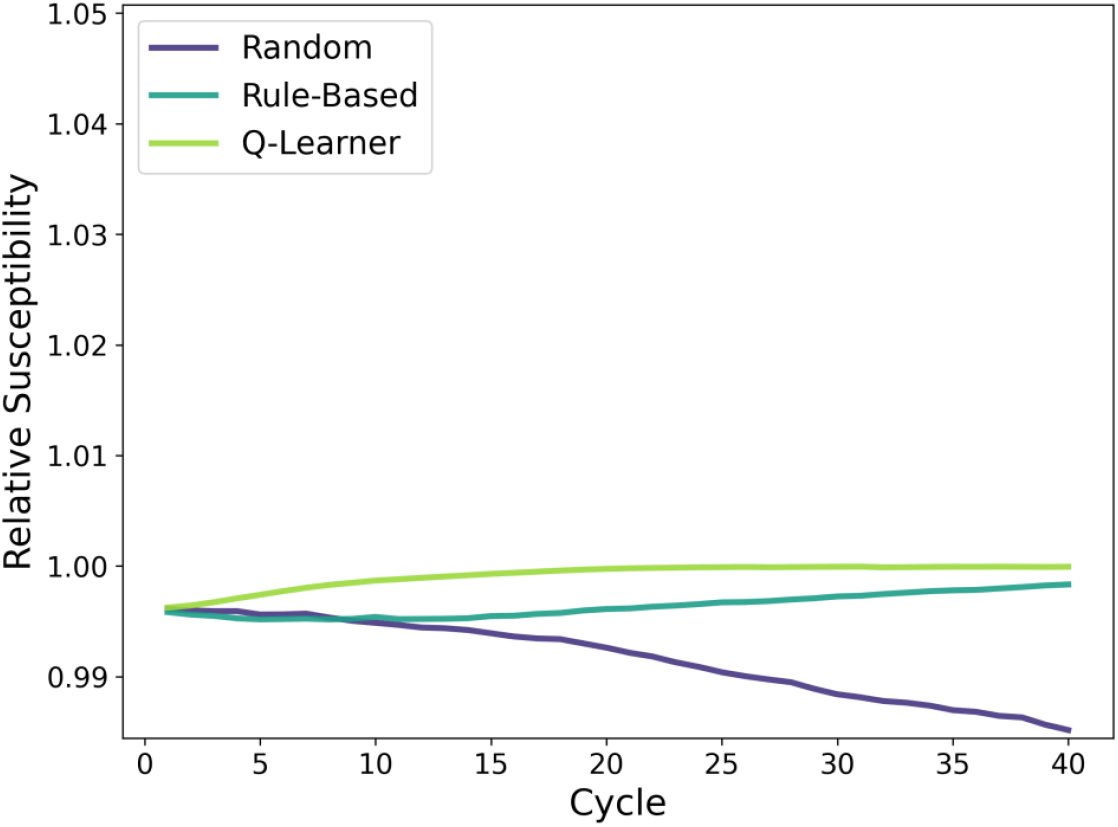
Resistance burden index across experimental cycles for the random, rule-based, and Q-learning agents.

These patterns were reflected in aggregate performance metrics across experimental runs. The Q-learning agent achieved lower mean AUC MIC values for both antibiotics and lower copper burden compared with the random and rule-based agents. In contrast, the random agent exhibited higher cumulative MIC exposure and copper burden, with greater variability across runs, while the rule-based agent demonstrated intermediate values across metrics Mean resistance burden index values remained near their upper bound across agents, indicating that between-agent differences primarily reflected divergence in resistance trajectories rather than short-term shifts in resistance burden.

This suggests that differences in AUC MIC primarily reflected divergence in resistance trajectories over time rather than short-term differences in growth suppression.

### 3.2. GLMM analysis of resistance trajectories

Results from generalized linear mixed models (GLMMs) aligned with the AUC-based summaries, providing model-based estimates of differences in simulated resistance trajectories across agents. Relative to the Q-learning agent, the random and rule-based agents were associated with higher MIC values over cycles for both antibiotics, with effect sizes varying by agent and antibiotic. Interaction terms between cycle and agent indicated differential temporal patterns in MIC trajectories. The random agent showed increasing MIC values over cycles relative to the Q-learning agent, while the rule-based agent exhibited smaller interaction effects, consistent with intermediate resistance trajectories. Across models, estimated effects for polymyxin B were generally smaller in magnitude than those observed for chloramphenicol, although the direction of effects was consistent across antibiotics.

## 4. Discussion

This study explored the application of reinforcement learning (RL) to model explore strategies for modulating antimicrobial resistance dynamics (AMR) under copper-driven antimicrobial pressure. Across simulation experiments, the Q-learning agent consistently achieved lower cumulative resistance exposure and copper burden compared to both rule-based and random policies, demonstrating the advantages of adaptive decision-making in sequential resistance management problems. These findings align with prior work framing AMR as a feedback-control problem, where adaptive policies can outperform static or heuristic strategies under noisy or delayed feedback conditions (*22,24*).

While the observed performance differences across agents were robust at the simulation level, this framework was intentionally designed as an exploratory model rather than a mechanistic representation of microbial evolution. As such, further refinement and empirical validation would be required to capture the full biological complexity of resistance dynamics observed in natural and clinical settings, including spatial structure, population heterogeneity, and ecological interactions (*17,31,32*). The present results should therefore be interpreted as illustrating relative agent behavior under controlled assumptions rather than predictive estimates of resistance outcomes.

This study establishes a baseline framework for applying reinforcement learning approaches to organic antimicrobial stressors, using copper as a representative compound with well-documented antimicrobial properties (*15–18*). In contrast to prior RL-based AMR studies that focus on synthetic antibiotics and empirically measured fitness landscapes (*22*), the present work emphasizes interpretability, limited state information, and simplified control actions. This design choice reflects scenarios in which detailed molecular diagnostics or high-resolution resistance measurements may be unavailable, such as resource-limited clinical or agricultural environments (*33,34*).

Several limitations of the current framework warrant explicit consideration. First, resistance evolution was modeled using stochastic update rules rather than explicit genetic or molecular mechanisms. While this abstraction is consistent with prior systems-level approaches to AMR modeling (*22,24*), it does not capture processes such as horizontal gene transfer, compensatory mutations, or multi-species interactions, all of which play critical roles in real-world resistance dynamics (*35,36*). Additionally, resistance burden was represented using an index derived from MIC values rather than empirical growth measurements or fitness assays, limiting direct biological interpretation of growth or viability outcomes. Third, copper was treated as a controllable environmental stressor rather than a therapeutic intervention, and the framework does not address toxicity, pharmacokinetics, or host-mediated effects that would be relevant in clinical translation (*17,37,38*). Despite these limitations, the framework provides a useful platform for examining how differing levels of adaptability influence resistance trajectories under antimicrobial pressure. The comparison between random, rule-based, and learning-based agents highlights how even simple adaptive policies can yield measurable improvements over static control strategies when resistance dynamics unfold over time. From a broader systems perspective, reinforcement learning offers a flexible methodological lens for studying AMR as a dynamic process shaped by feedback between microbial populations, antimicrobial exposure, and environmental constraints (*22,24,33,34,39*).

Future work could extend this framework by incorporating additional sources of biological complexity, including multi-species populations, spatial structure, and horizontal gene transfer mechanisms (*35,36*). Further studies could also evaluate alternative organic antimicrobial compounds or combinations thereof, as well as more advanced learning algorithms capable of handling continuous state spaces and delayed rewards. Rather than providing a definitive solution to antimicrobial resistance, this study contributes an interpretable and extensible modeling framework that bridges microbiology and adaptive decision-making, laying groundwork for future investigations into AI-assisted AMR control strategies.

## 5. Conclusions

This study presented an exploratory reinforcement learning (RL) framework for modeling antimicrobial resistance (AMR) dynamics under copper-driven antimicrobial pressure. By comparing a random policy, a rule-based heuristic, and a Q-learning agent within a controlled simulation environment, this work demonstrates how differing levels of adaptability influence resistance trajectories over time. Across experiments, the Q-learning agent consistently achieved lower cumulative resistance exposure and copper burden than non-adaptive or fixed-decision strategies, illustrating the potential advantages of adaptive control in sequential resistance management problems.

Importantly, the framework was designed to emphasize interpretability and conceptual clarity rather than biological or computational complexity. Resistance evolution and resistance burden were modeled using simplified, stochastic dynamics grounded in commonly reported microbiological indicators, such as minimum inhibitory concentration (MIC), without explicitly modeling molecular mechanisms. This design choice reflects scenarios in which detailed genetic data or high-resolution diagnostics may be unavailable, while still allowing meaningful comparison of control strategies under antimicrobial pressure.

Rather than proposing copper as a therapeutic intervention, this study uses copper as a representative organic antimicrobial stressor to explore broader methodological questions at the intersection of microbiology and adaptive decision-making. The results highlight how even simple reinforcement learning approaches can outperform static or heuristic policies when resistance dynamics unfold across repeated exposure cycles. These findings support the use of RL as a flexible modeling tool for studying AMR as a dynamic, feedback-driven process. The value of this framework lies not in biological realism, but in providing a transparent testbed for comparing decision strategies under shared and explicitly constrained assumptions.

The framework introduced here is intentionally modular and extensible. Future work may incorporate additional biological complexity, such as multi-species interactions, spatial structure, or alternative antimicrobial compounds, as well as more advanced learning algorithms capable of operating in continuous or partially observed environments. In parallel, empirical validation and tighter integration with experimental data would be necessary steps toward translational relevance.

Taken together, this work introduces a transparent and extensible simulation framework integrating microbiological indicators with reinforcement learning, establishing a foundation for future methodological research on antimicrobial resistance dynamics under constrained or uncertain conditions.

## Supporting information

https://github.com/h-hedman/healthcare-data-science/tree/main/organic-amr-rl

## Supplementary Materials

All relevant code, data, and figures are available in SM1.

## Disclaimer/Publisher’s Note

The statements, opinions and data contained in all publications are solely those of the individual author(s) and contributor(s) and not of MDPI and/or the editor(s). MDPI and/or the editor(s) disclaim responsibility for any injury to people or property resulting from any ideas, methods, instructions or products referred to in the content.

