## Supplementary material for "A Reinforcement Learning Approach for Modeling Organic Compound-Induced Antimicrobial Resistance Dynamics": https://github.com/h-hedman/healthcare-data-science/tree/main/organic-amr-rl

Supplemental Materials 1 – Code

The code used for this study is available in the public repository at: https://github.com/h-hedman/healthcare-data-science/tree/main/organic-amr-rl.

All scripts required to reproduce the models, tables, and figures are provided.
